# Temporal dynamics of microbial transcription in wetted hyperarid desert soils

**DOI:** 10.1101/2021.05.12.443739

**Authors:** Carlos León-Sobrino, Jean-Baptiste Ramond, Clément Coclet, Ritha-Meriam Kapitango, Gillian Maggs-Kölling, Don A. Cowan

## Abstract

Rainfall is rare in hyperarid deserts but, when it occurs, it triggers large biological responses which are considered to be essential for the long-term maintenance of biodiversity. In such environments, microbial communities have major roles in nutrient cycling, but their functional responses to short-lived resource opportunities are poorly understood. We used whole community metatranscriptomic data to demonstrate structured and sequential functional responses in desiccated desert soils to a simulated rainfall event over a seven-day time frame. Rapid transcriptional activation of Actinobacteria, Alpha-proteobacteria and phage transcripts was followed by a marked increase in protist and myxobacterial activity, before returning to the original state. In functional terms, motility systems were activated in the early phases, whereas competition-toxicity systems increased in parallel to the activity from predators and the drying of soils. The dispersal-predation dynamic was identified as a central driver of microbial community responses to watering. Carbon fixation mechanisms that were active under dry condition were rapidly down-regulated in wetted soils, and only reactivated on a return to a near-dry state, suggesting a reciprocal balance between carbon fixation and fixed-carbon acquisition processes. Water addition induced a general reduction in the transcription of stress response genes, most prominently HSP20, indicating that this chaperone is particularly important for life in desiccated ecosystems. Based on these data, we propose a rainfall response model for desert soil microbiomes.

## Introduction

Arid lands cover approximately one third of the terrestrial surface (Laity, 2009). Aridity, as a result of very low precipitation and high potential evapotranspiration rates, severely limits biomass production and promotes oligotrophy (Delgado-Baquerizo et al., 2013; Maestre et al., 2015), eventually leading to habitat fragmentation where productivity is concentrated in sheltered “islands” (Pointing and Belnap, 2012; Schlesinger et al., 1995). Microorganisms in open soils outside of these privileged micro-environments are subjected to an intense desiccation stress which limits their biological activity (Lebre et al., 2017). However, desert edaphic microorganisms are regarded as important for the long-term fertility of soils and for post-rainfall grass germination (Delgado-Baquerizo et al., 2016). The Namib Desert, on the south-western coast of Africa, is the oldest continuously hyperarid desert in the world and provides an excellent model of a stable hyperarid ecosystem (Seely et al., 2008). It is generally accepted that much of the microbiome in such environments exists in a state of dormancy (Bär et al., 2002; Lebre et al., 2017), but responds with dramatic changes in structure and activity when water becomes available (Armstrong et al., 2016; Austin et al., 2004; Collins et al., 2014; Garcia-Pichel and Pringault, 2001; Štovíček et al., 2017). Recent studies have, however, shown that some transcriptional activity is detectable in hyperarid soil microbial communities during prolonged dry periods (Gunnigle et al., 2017; León-Sobrino et al., 2019; Schulze-Makuch et al., 2018). Since arid lands represent a large part of the terrestrial surface of the Earth and are currently expanding through desertification processes (Huang et al., 2016), a deeper understanding of these pulsed phenomena is of global relevance.

Research on desert soil microbial ecology has primarily focused on bacterial communities, suggesting that extreme abiotic pressures, such as high temperature, desiccation stress and UV radiation are dominant drivers of both the diversity and function of bacterial communities (Frossard et al., 2015; Johnson et al., 2017; León-Sobrino et al., 2019; Mandakovic et al., 2018; McHugh et al., 2017; Ronca et al., 2015; Scola et al., 2018; Vikram et al., 2016). By comparison, virus/phage communities in hot desert soils have been relatively under-studied (Trubl et al., 2020; Williamson et al., 2017), despite the fact that they represent one of the most abundant and genetically diverse entities on Earth and are inferred to play central ecological roles in biogeochemical nutrient cycling in diverse environments (Adriaenssens et al., 2017; Bezuidt et al., 2020; Luo et al., 2020; Zablocki et al., 2018). Furthermore, phages (i.e., viruses that infect bacteria) have been shown to affect community turnover and resource availability in soils and to globally drive microbial evolution via horizontal gene transfer (Rodriguez-Valera et al., 2009; Trubl et al., 2019; Williamson et al., 2017).

In this study, a gravel plain soil plot in the central Namib Desert was subject to an artificial rainfall pulse of 30 L/m^2^, the approximate average annual rain received in this region (25 mm) (Eckardt et al., 2013) and sufficient to stimulate plant germination (Seely et al., 2008). The short-term responses of the soil microbiome were monitored using whole-transcriptome analysis of gene function. Based on this analysis, we propose a structured water response model with differentiated phases and trophic interactions.

## Materials and methods

Surface soils were collected from two adjacent 3.5 × 3.5 m plots (approx. 10m separation) in the central Namib Desert gravel plains (23° 33’ 18’’ S, 15° 3’ 20’’ E). The control plot remained dry, while the experimental plot was manually watered (30L/m^2^, simulating a 30 mm rainfall event) at T0 (26th April 2017 10:00 AM WAT/UTC+1), using a synthetic “Namib Desert rain” solution, prepared from ultrapure DNA-free water supplemented with a defined salt mixture (Frossard et al., 2015). Plots were subdivided into 0.5 × 0.5 m quadrats and randomly sampled, in triplicate, at specified times after water addition (Supplementary Figure S1). 20 g samples of surface (0-5 cm) soil were collected at 10 minutes; 1, 3 and 7 hours; and 1, 3 and 7 days after simulated rainfall, preserved immediately in RNAlater solution (Sigma-Aldrich, St. Louis MO, USA) and subsequently frozen at -20°C prior to RNA extraction. Additional 200 g soil samples were collected from the same locations into WhirlPak™ bags (Nasco, Fort Atkinson WI, USA) and frozen for subsequent soil chemistry analysis.

The water content of soil samples (sieved to <2 mm particles) was measured gravimetrically for three days after watering. Soil silt, sand, and clay compositions were measured by the hydrometer method (Gee and Bauder, 1986). Particle size distributions were determined by sieve separation (> 1000 µm, > 500, > 250, > 100, > 53 and < 53 µm). pH, electrical conductivity, Na, Cl, K, Ca, Mg, NO3, NH4 and P composition were analysed at Bemlab (Pty) Ltd. (Strand, Western Cape, South Africa) using standard protocols. Soil organic carbon percentage was measured using the Walkley-Black test (Walkley, 1935).

RNA was extracted following the protocols described in León-Sobrino *et al*. 2019 (León-Sobrino et al., 2019) from triplicate soil samples. In order to mitigate the effect of chemical variation, the two biological replicates most similar to the average chemical composition of all sampled soils were selected for RNA extraction. Stranded, rRNA-depleted libraries were prepared with the ScriptSeq Complete Gold Kit (Epidemiology) (Illumina, San Diego, USA) and 150 bp paired–end sequences were read on a HiSeq4000 platform (Illumina).

Sequencing outputs were processed using the BBtools suite v. 38.26 (Bushnell et al., 2017) (https://sourceforge.net/projects/bbmap/). Read ends below a quality Phred value of 20 were trimmed using *BBDuk*; rRNA and human RNA sequences were identified and removed using SILVA v. 111 (July 2012) (Quast et al., 2012) and 5SRNA (Szymanski et al., 2016) databases and a curated human genome reference assembly hG19 (https://drive.google.com/file/d/0B3llHR93L14wd0pSSnFULUlhcUk) (bbmap, 2018) following recommended protocols. Optical duplicates generated by the patterned sequencing flowcell were removed using the *Clumpify* function from the BBtools suite, setting the distance cut-off to 2,500 pixels. Transcript assembly was performed using transAbyss v.2.0.1 (Robertson et al., 2010) for each library. Individual assemblies were merged with the same software (*transabyss-merge* function) to generate a reference metatranscriptome.

Contigs were annotated at the Integrated Microbial Genomes and Microbiomes (IMG/M) server (Huntemann et al., 2015) (https://img.jgi.doe.gov/). Predicted protein products from genes were also analysed against the Conserved Domain Database (CDD) using Delta Blast (e-value threshold 10^−4^) (Boratyn et al., 2012; Marchler-Bauer et al., 2015). Contig taxonomy was determined by a consensus among all individually classified genes, requiring a *quorum* of >50% at each taxonomic level.

Reads were aligned to the reference assembly using BBmap (Bushnell et al., 2017) and reads for annotated regions in each library were counted using FeatureCounts v. 1.6.3 (Liao et al., 2014). The assembled counts matrix was aggregated along functional and/or taxonomic categories as required for each analysis.

Differential transcription along the time series was analysed with the DESeq2 v. 1.14 package in R (Love et al., 2014), comparing data from the control (dry) soil samples with those from the experimental (wetted) samples from the same point in the time-series. Transcripts that significantly diverged in abundance from the control at any given time were considered up-regulated in response to watering (adjusted p-value ≤ 0.05 for the likelihood ratio test). Transcripts per million (TPM) of ribosomal protein genes as a fraction of the total for their respective taxonomic group (Rp:T) were employed to estimate absolute activity of each group along the time course, following the premise that ribosome densities in a cell relate to metabolic activity and growth rates (Bosdriesz et al., 2015; Bremer and Dennis, 1996).

Viral contigs were identified from assembled transcriptomes using VirSorter v. 1.0.3 (Roux et al., 2015) and the virome database on the iVirus platform hosted by CyVerse (Bolduc et al., 2017). Only contigs >1 kb, and classified as categories 1, 2, 4, and 5 were considered (phages and prophages identified with the “pretty sure” and “quite sure” qualification). To calculate the relative abundances of the different viral contigs in each transcriptome, quality filtered metatranscriptomic reads were mapped back to the viral contigs with Bowtie2 v. 2.2.6, using the default parameters (Langmead and Salzberg, 2012). The output SAM files were converted into BAM files, sorted and indexed, using SAMtools (Li et al., 2009). A python script was used to generate a coverage profile of viral contig abundances across samples.

ORFs were predicted within putative viral contigs using Prodigal (Hyatt et al., 2010). Transcripts per million (TPM) putative viral protein genes were employed to normalize the final ‘viral OTU’ (vOTU) values for each sample. Predicted protein sequences were clustered with proteins from viruses in the NCBI ViralRefSeq-prokaryotes-v85 based on all-versus-all BLASTp search with an E value of 1 × 10^−3^, and clusters were defined with the Markov clustering algorithm and processed using vConTACT2 (Bin Jang et al., 2019). The stringency of the similarity score was evaluated through 1,000 randomizations by permuting protein clusters or singletons (proteins without significant shared similarity to other protein sequences) within pairs of sequences having a significance score ≤ 1 (negative control). Subsequently, pairs of sequences with a similarity score > 1 were clustered into viral clusters with the Markov clustering algorithm using an inflation value of 2. The resulting gene-sharing network from vConTACT2 classification was visualized with Cytoscape software v. 3.7.0 (Smoot et al., 2011). Reference sequences from RefSeq database that co-clustered with the putative viral sequences were used to predict viral taxonomy.

## Results

### Site characteristics and taxonomic analysis

The sample site (Figure 1A and 1B) was a characteristic Namib Desert calcrete gravel plain (Gombeer et al., 2015) with high sand composition (92 ± 1.6 %) and very low organic carbon content (0.04%) (Supplementary Table S1). Soil chemical composition was relatively homogeneous in all sampled sectors and between the sample and control sites (Supplementary Table S1). Gravimetric water content measurements showed that more than half of the water content in the surface soil (0-5 cm) was lost 24 hours after the simulated 30 mm rainfall. After 3 days, the water content was similar to that of the dry control (Figure 1C).

**Figure 1:**
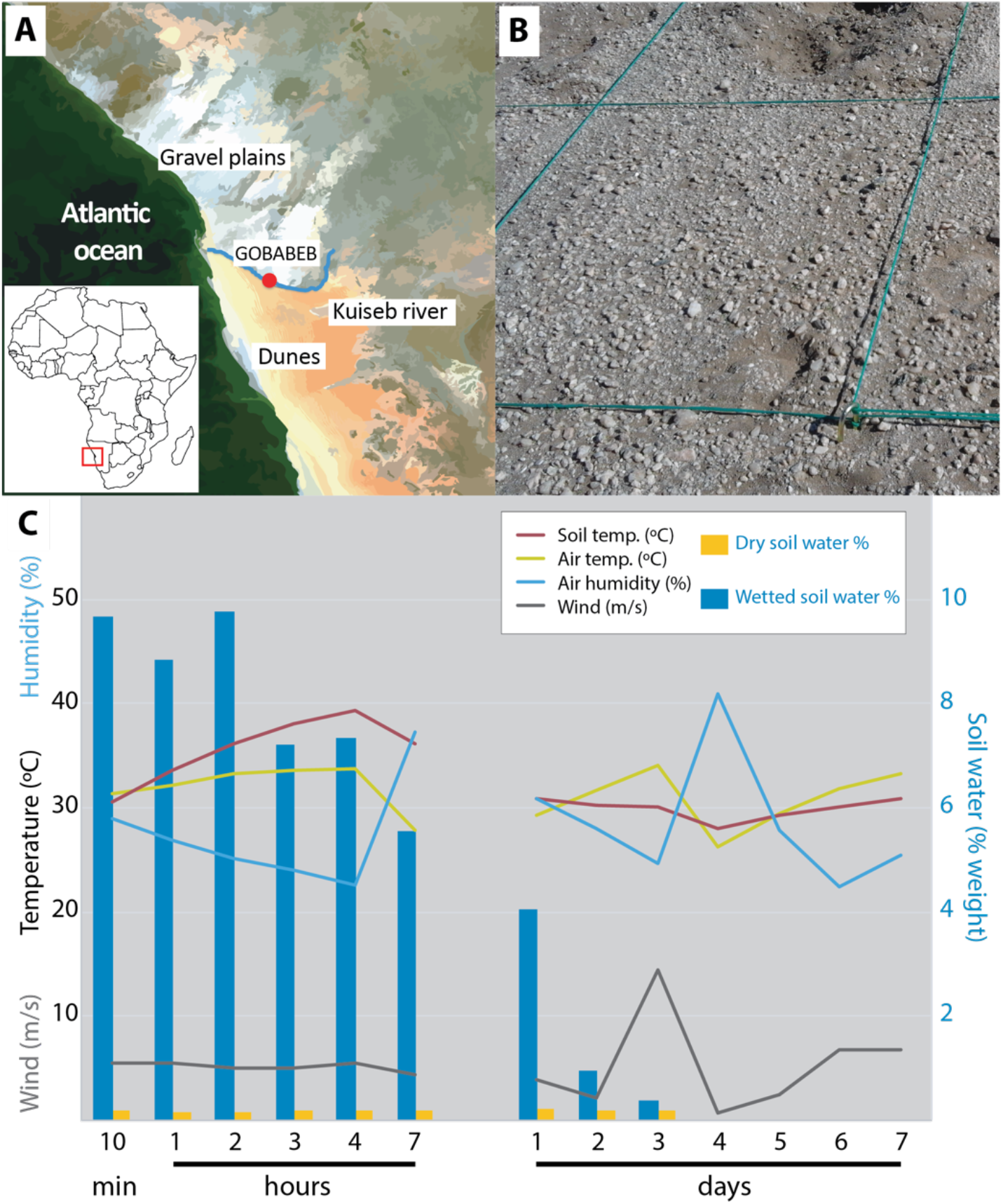
Sample site and environmental conditions of the sampled soils. A) Map of the Namib Desert gravel plain location. Modified from European Space Agency, ESA/Envisat <u>CC BY-SA 3.0 IGO</u>. B) View of a representative portion of the gravel plain sampling site. C) Environmental conditions over the experimental period: Air temperature, humidity and wind were recorded by the nearby Gobabeb meteorological station (Southern African Science Service Centre for Climate Change and Adaptive Land Management (SASSCAL), station 8893).

Transcript read assembly yielded a consensus metatranscriptome of 208.95 Mb in which 378 802 coding regions were annotated, including 372 044 predicted protein-coding genes. On average, for the 24 sequenced libraries, 61.7% of reads could be aligned back to contigs. Functions were predicted for 29.9% of the protein-coding genes using the KEGG database (Kanehisa et al., 2016), and 51.6% using the CDD. 56.8% of contigs were taxonomically classified at phylum level, and 56.3% down to family level.

A functional and taxonomic analysis of the transcriptional profiles revealed large differences in gene expression levels between treatment and control soil transcriptomes within 10 minutes after watering (Figure 2, Supplementary Figure S2). Communities from dry soils were characterized by stable (i.e., largely unchanged) transcription over the 7-day experimental period. In contrast, microbial communities in the watered soil plot underwent an immediate and large change in gene expression (Figure 2A) that progressively returned to basal expression levels within 7 days from the wetting event. Taxonomy-only transcriptional profiles followed this cyclical pattern (Figure 2B) but were less robust than KO gene categories in representing the changes in the soil microbiome induced by soil wetting.

**Figure 2:**
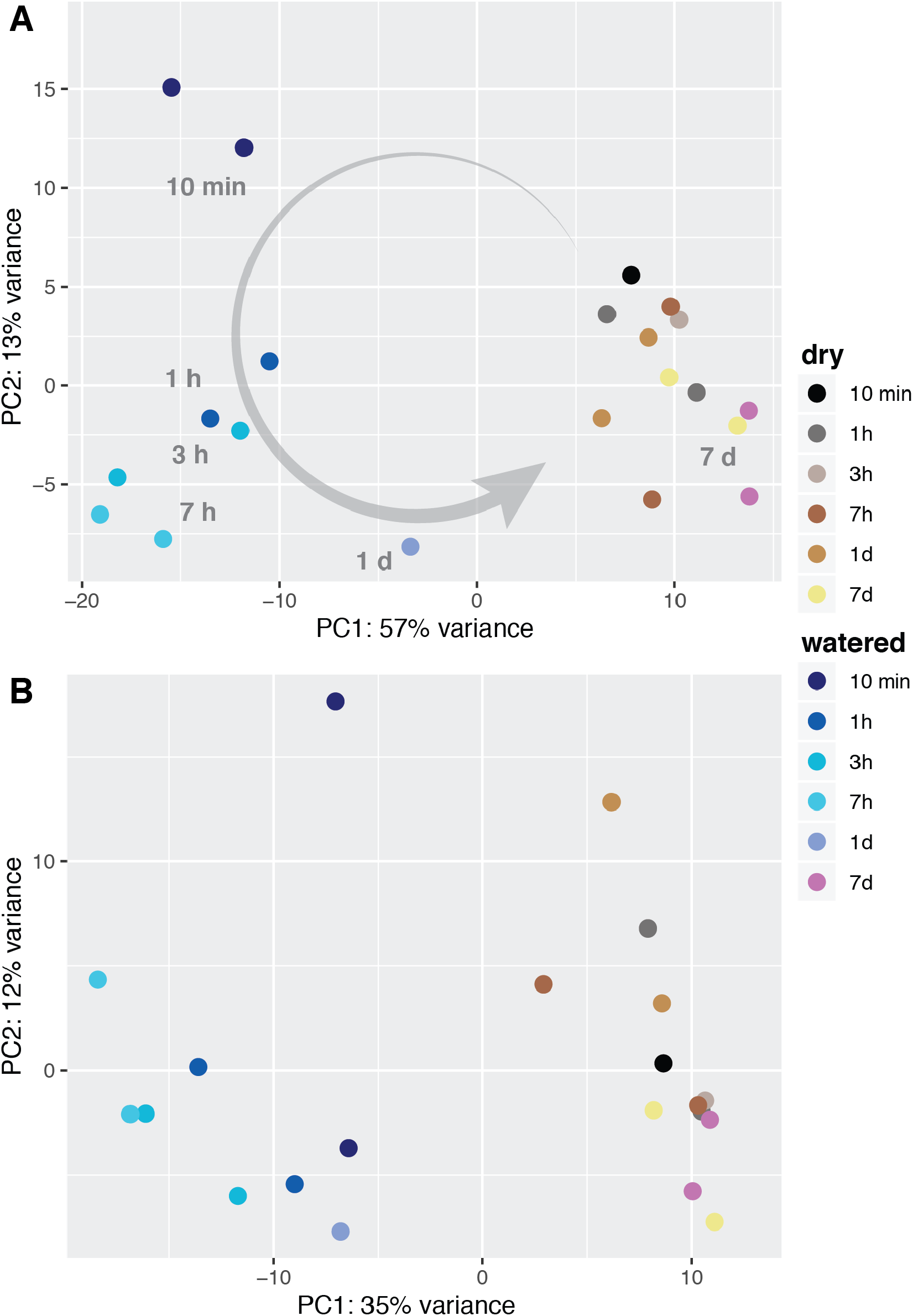
Principal components analysis of A) KEGG Orthologs (KO) annotated transcripts, and B) taxonomic (family) assignment of transcripts. The arrow designates the temporal transition pathway.

In both watered and dry soils, Actinobacteria and Proteobacteria were the most abundant phyla (average 40% and 16% in all samples, respectively, see Supplementary Table S2). Strikingly, transcripts from Nitrososphaeria, a class of ammonia oxidizing Thaumarchaeota, were the third most abundant (average 14% of classified TPM in all samples), suggesting that these taxa play an important role in the microbial community and in local nitrogen cycling. Protist transcripts (Oligohymenophorea class and Dictyostellales), which were rare in dry soil samples (<0.5%), increased to >4% within 7 hours of soil watering. Conversely, transcripts of fungal origin were more common in dry soils (1% of classified TPM) than in watered ones.

The production of ribosomes in each taxonomic group was analysed, as an indicator of their cellular metabolic activity, by quantifying ribosomal protein transcripts as a fraction of the total (Rp:T) (Bosdriesz et al., 2015; Bremer and Dennis, 1996). Ribosomal protein transcripts constituted a stable proportion for each taxon in dry control samples, typically <4% TPM (Figure 3). After water addition, however, Rp:T increased for all major taxa, peaking at characteristic times over the course of the experiment and returning to basal values by the end of the 7 day period. Actinobacteria, Alpha-proteobacteria and Chloroflexi were examples of ‘early-activation’ groups, with Rp:T values peaking within the first hour after water addition. Gemmatimonadetes experienced the largest relative increase in Rp:T, despite being a minor component of the soil microbiome (<1% TPM). A delayed rise in activity, peaking at 7 hours after the simulated rainfall, was evident for Delta-proteobacteria, protists and Bacteroidetes, especially those belonging to the Cytophagia class, which constituted the principal ‘late-activation’ group. The Pezizomycetes, the most active fungal class, also showed a slower response to water addition. Intermediate response patterns, with maximal Rp:T values at 3 hours after water addition, were observed for taxa such as Planctomycetes. Thaumarchaeal Rp:T ratios showed only moderate changes after watering and a relatively even distribution throughout the experiment.

**Figure 3:**
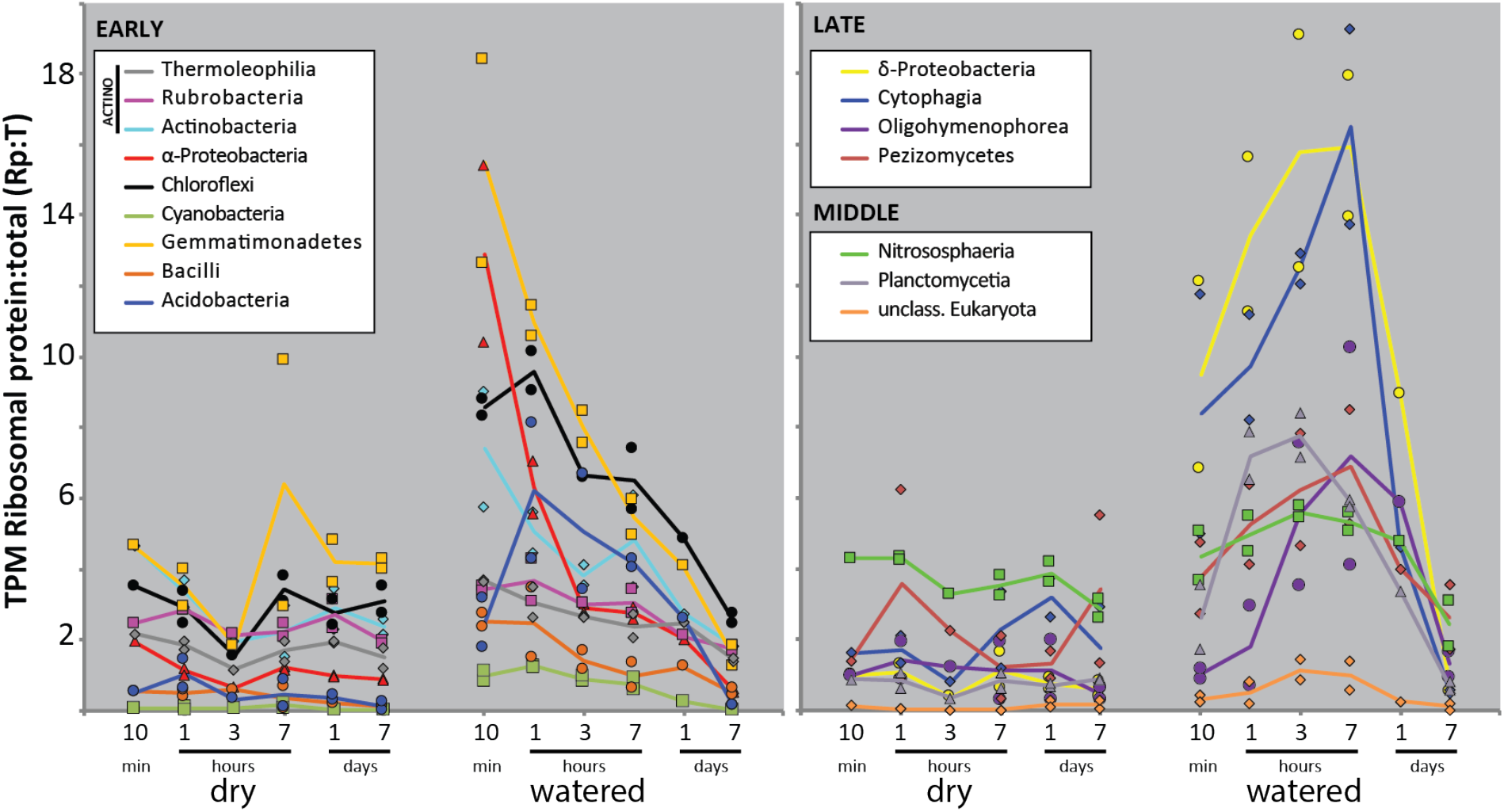
Ribosomal protein gene transcripts as a fraction of the total (Rp:T) among the most transcriptionally active microbial classes in the Namib soil community (>1% TPM average in dry or watered samples). Left panel shows *early* response classes whose Rp:T ratios peak within the first hour after watering. Right panel includes classes with Rp:T maxima 3 and 7 hours after watering (*middle* and *late* response taxa, respectively).

### Temporal changes in core cellular functions

Watering induced a general reduction of stress resistance gene transcripts in the microbial community immediately after watering (e.g., *groEL, dnaK, clp, tre*). Trehalose synthesis genes (*tre*), which drive solute accumulation under conditions of water stress (Lebre et al., 2017), declined in relative transcription in watered soils. Transcripts for the major chaperones *groEL* and *dnaK*, and the *clp* protease involved in protein misfolding control were also reduced. The most conspicuous change in core stress resistance systems was, however, that of the heat-shock protein HSP20 genes (KO13993, CD223149 or CD278440). Genes for this ATP-independent chaperone experienced the largest and most consistent transcript reduction across all significant (average ≥1 % of transcripts) bacterial taxa after watering (Figure 4), suggesting that HSP20 represents an archetypal “desiccation adaptation” gene.

**Figure 4:**
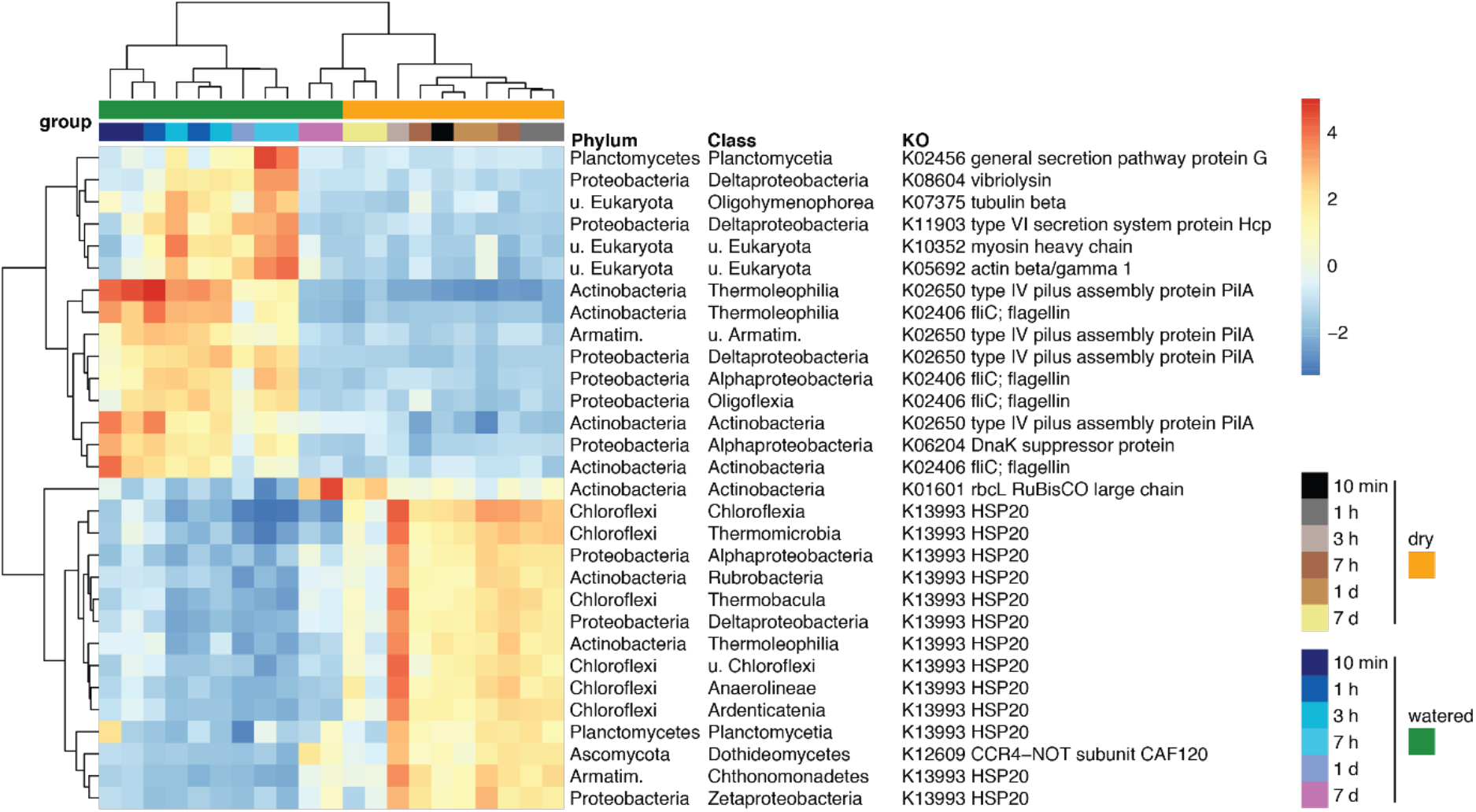
Temporal transcription patterns of the 30 most variable genes among those with significantly differential transcription along the time series. Transcript data was aggregated along taxonomic (class) as well as functional (KEGG Orthologs) groups for the analysis. Values were normalized using the Variance Stabilizing Transformation (*DESeq* R package).

Motility related transcripts were rapidly affected by soil wetting (from 10 minutes after water addition). Flagellar genes were up-regulated within the first hour in the dominant Actinobacteria and Alpha-proteobacteria (Figure 4) and in less dominant taxa such as Planctomycetes and Firmicutes. Type IV pilus genes, involved in gliding motility, were also rapidly upregulated after water addition, most notably in Actinobacteria. These responses suggest that rapid dispersal is a key behavioural response to soil wetting, particularly in Actinobacteria.

A significant up-regulation of Eukaryotic adhesion and cytoskeletal components, myosin, actin and tubulin, was evident 3 to 7 hours after soil wetting (Figure 4), but was limited to non-fungal taxa, particularly in Oligohymenophorea (ciliate) and Dictyosteliales (slime mould).

Another highly significant increase in transcriptional activity observed several hours after water addition involved interbacterial competition and predation genes. These included type VI secretion systems (T6SS) and vibriolysin encoding genes from the order Myxococcales (Delta-proteobacteria), and T6SS and serralysin encoding genes from Alpha-proteobacteria. Heightened T6SS transcription was also observed in several other bacterial taxa, such as Planctomycetes, Gemmatimonadetes and Gamma-proteobacteria, suggesting that antagonistic interactions are of significant ecological importance in newly wetted soils.

### Biogeochemical cycles

#### Carbon utilization

Despite the observation that microbial photosynthetic processes are water-limited in hyperarid soils (Tracy et al., 2010; Warren-Rhodes et al., 2006), transcript data did not show a significant increase in cyanobacterial gene expression after water addition, and both overall cyanobacterial transcript abundance and Rp:T ratios remained low (Figure 3, Supplementary Table S2). Furthermore, core carbon fixation genes, including RuBisCO and carboxylases genes from chemoautotrophic pathways, consistently showed reduced relative transcription in wetted soils. Thaumarchaeal carbon fixation was seemingly also affected by watering, reflected in the transient inhibition of 4-hydroxybutyryl-CoA dehydratase from the 3-hydroxypropionate/4-hydroxybutyrate cycle following inundation. These data are indicative of an inhibition of both photosynthetic and non-photosynthetic carbon fixation mechanisms immediately after soil saturation (Fig S3A). However, expression of carbon fixation genes increased by the end of the 7-day experimental period, when soil water content had returned to basal levels (Figure 4, Supplementary Figure S3A).

Dramatic increases in CO2 emissions from newly wetted desert soils, attributed to degradation of dissolved organic matter, have been widely reported (Austin et al., 2004). Soil chemistry, however, showed no significant reduction in soil Total Organic Carbon (Supplementary Table S1), possibly due to limed sensitivity of the employed method at the very low values present in these soils. In our transcript data, indicators of biomass degradation, including carbohydrate and peptide transport systems (e.g., *rbsB, xylF*, and *livK, dppA*) significantly increased for most taxa immediately after water addition. The Bacteroidetes phylum, dominated by the Cytophagales, showed dramatic up-regulation of subtilisin protease transcripts and components of the protein and carbohydrate import machinery (*ragA*/*susC* CD274948; *susD* CD185760), suggesting an important role of this group in the turnover of organic matter in the ecosystem.

#### Nitrogen and phosphate utilization

Transcript data suggested that Thaumarchaea were the main drivers of nitrogen cycling in dry soils. Ammonia monooxygenase and NO-forming nitrite reductase (*amo* and *nirK*) transcripts, originating largely from Thaumarchaea, were among the most abundant overall. Transcripts associated with both processes declined after soil watering (Figure 5A, Supplementary Figure S3B). All other detected transcripts implicated in ammonification, such as nitrate and nitrite reductases *narGH* and *nirAD* (mainly transcribed by Nitrospirae and Actinobacteria, respectively), and nitric oxide reductase (*norC*), were also reduced after watering. Conversely, peptide transporter transcripts (*livKH, dppA*) significantly increased in response to water addition for several taxa (Actinobacteria, Alpha-, Beta- and Delta-proteobacteria), suggesting that organic nitrogen dominates N acquisition processes in newly wetted soils.

**Figure 5:**
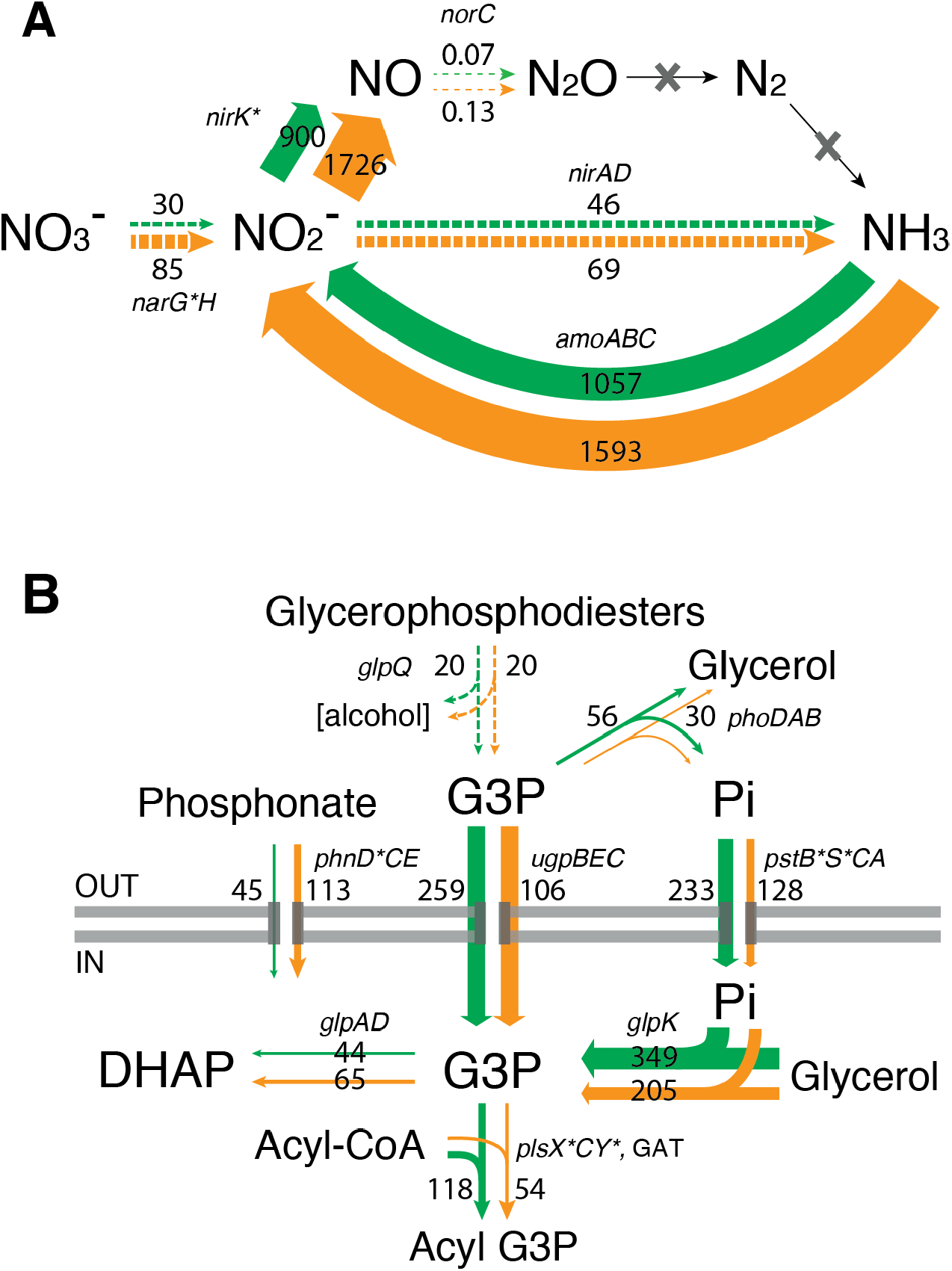
A) Nitrogen cycling and B) phosphate assimilation system transcription in the surface soil microbial community. Average TPM measurements are provided for the dry plot samples (orange arrows) and wetted samples up to 7 hours post watering (green arrows). Asterisks indicate significant differentially transcribed KOs at community level at any point along the time series. Panel B figure modified from León-Sobrino *et al*., 2019.

Changes in phosphorus acquisition pathways were also identified. Transcript data suggested that phosphate acquisition after water addition was dominated by the proteobacterial *ugpB sn*-glycerol 3-phosphate (G3P) transport system subunit, particularly in Alpha-proteobacteria, followed by the *pst* inorganic phosphate transporter (Figure 5B). By contrast, phosphonate transport (*phn*) gene transcription was reduced after soil wetting.

### Analysis of viral transcripts

A total of 68 contigs were characterized as being of viral origin using VirSorter (Supplementary Figure S4). In both dry and watered soil samples, viral contigs were significantly lower than bacterial contigs (ANOVA, *p* < 0.01), representing an average of 0.12 ± 0.07% TPM of viral protein transcripts. Over the period of the experiment, viral reads exhibited a significant temporal pattern, where read numbers remained low in all dry soil samples but exhibited a bimodal response in wetted soils (Figure 6). An initial very rapid increase in viral RNAs (approx. 2.2-fold at the first post-watering sample point: 10 min) was followed by a 12-fold decline over 7 hours. A second increase of viral read numbers (approx. 6-fold) occurred between 7 hours and 7 days.

**Figure 6.**
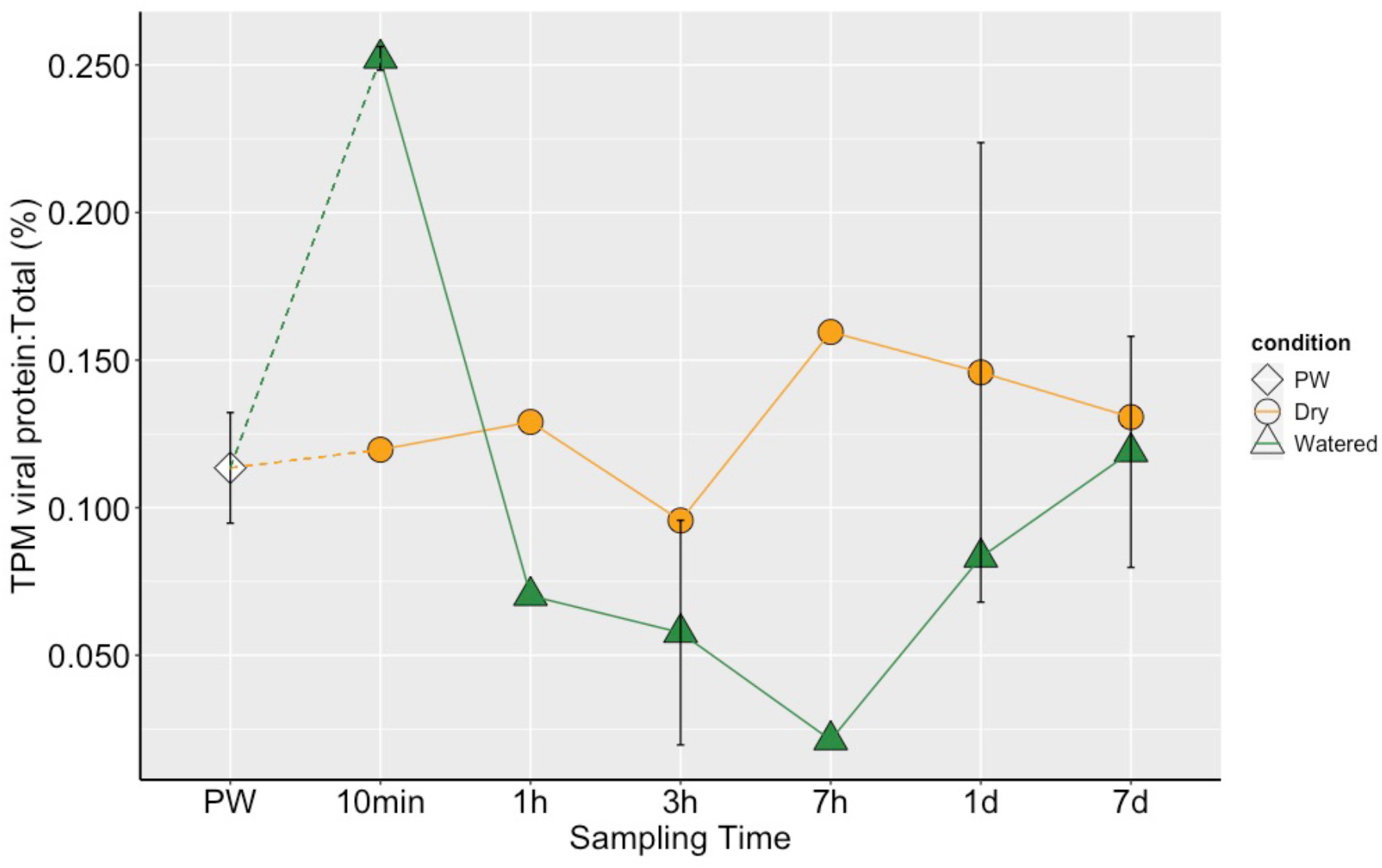
Temporal changes in viral protein gene transcripts as a fraction of the total (Vp:T ratios; %) for watered and dry (control) soil samples. The PW (Pre-watering) value represents the mean ± sd of Vp:T ratios in all control samples.

To investigate the diversity of the ‘active’ viral population, we used a genome-based network analysis of the shared protein content with the prokaryotic viral genomes (RefSeq v85). This analysis grouped 35 viral contigs into viral clusters (VCs) while 33 viral contigs remained non-clustered and were considered as singletons (Figure 7). In the network, 10 VCs containing viral contigs from our study were predicted, 7 of which did not belong to VCs with RefSeq virus genomes but instead clustered together into novel VCs, and 3 of which could be assigned taxonomy at the family level (Figure 7) as members of the *Caudovirales* (*Siphoviridae* and *Leviviridae*).

**Figure 7.**
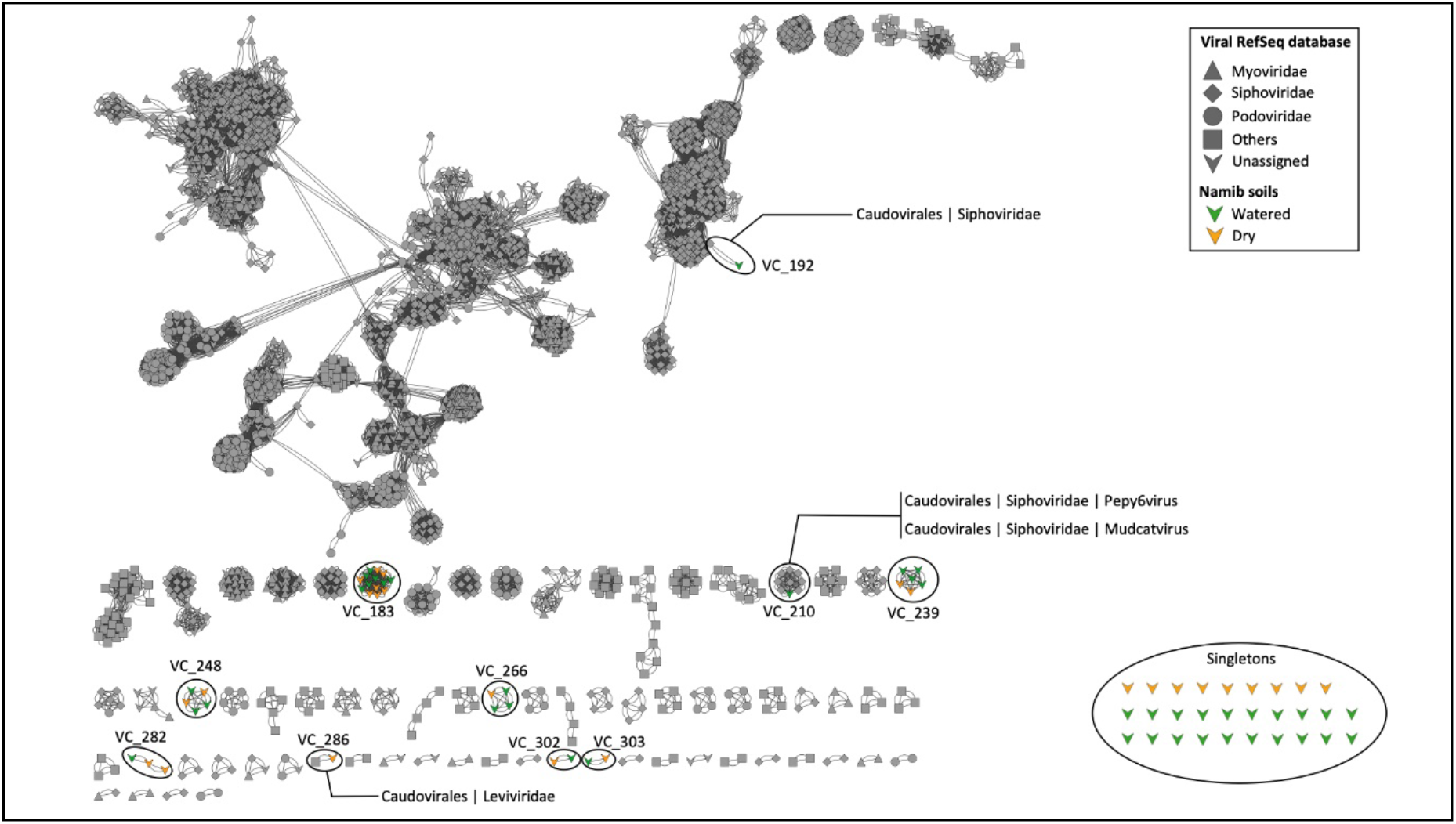
Network analysis, relating Water Experiment (WE) phage sequences to known viral sequences from the RefSeq Database. Circled clusters present viral contigs identified in this study. Shapes indicate major viral families, RefSeq sequences are in grey and WE contigs are colored (yellow and green for dry and watered samples, respectively). Each node is depicted as a different shape, representing viruses belonging to Myoviridae (triangle), Podoviridae (circle), Siphoviridae (diamond), or uncharacterized viruses (V shape). Single shapes represent viral singletons identified in this study. Edges (lines) between nodes indicate statistically weighted pairwise similarity scores (see Materials and Methods) of ≥ 1.

## Discussion

Studies of the microbial ecology of desert edaphic niches have tended to focus on biological ‘hotspots’: hypoliths, soil crusts, and soils in the immediate vicinity of plants (e.g., Pointing and Belnap, 2012; Ramond et al., 2008; Marasco et al., 2018). While the use of high-throughput sequencing techniques (Crits-Christoph et al., 2013; Fierer et al., 2012, 2007; León-Sobrino et al., 2019; Jordaan et al., 2020; Vikram et al., 2016) has greatly expanded of knowledge of the microbial ecology of these niches, most studies have centred on microbial diversity rather than functionality. Total RNA sequencing is therefore a valuable tool to monitor microbial functionality at a high temporal resolution, particularly since mRNA is only generated by active organisms and is ephemeral, leaving little to no legacy signal (León-Sobrino et al., 2019; Rajeev et al., 2013; Steven et al., 2018).

Previous studies have clearly demonstrated a long-term compositional and functional adaptation of Namib Desert edaphic and hypolithic microbial communities to abiotic factors, particularly water input regimes histories (e.g., fog *vs* rain inputs; Cowan et al, 2020; Frossard et al., 2015; Ramond et al., 2018; Scola et al., 2018; Stomeo et al., 2013;). The location for this study site was situated in the central hyper-arid zone of the Namib Desert, which receives little input water from either rain or fog (Eckardt et al., 2013). The location should therefore optimise the responsiveness of the soil microbiome to water input (Seely et al., 2008).

### The HSP20 chaperone is important for microbial life in desiccated soils

One of the projected effects of water addition was an apparent reduction in cellular stress, suggested by the immediate down-regulation in transcription of stress-resistance genes. The most conspicuous of these changes was the abrupt reduction in transcription - across all major bacterial taxa - of the small ATP-independent heat-shock protein HSP20. This chaperone has been characterized as a broad-spectrum bacterial stress resistance mechanism (Bepperling et al., 2012; Haslbeck and Vierling, 2015).

### Desert soil microbial community members are sequentially activated after a water event

Data from the control (unwatered) site confirmed the presence of a diverse and functionally active microbial community in desiccated desert soils (see also, Gunnigle et al., 2017; Jordaan et al., 2020; León-Sobrino et al., 2019; Schulze-Makuch et al., 2018). However, out data suggest a level of remarkable “metabolic readiness”, with a dramatic increase in transcription associated with previously inactive taxa occurred within 10 minutes after water addition. We note that transcriptional response rates may be considerably faster, given that 10 minutes was the first sampling time-point. In polyextreme hyperarid desert soils, where most (micro)organisms remain in a state of metabolic dormancy, such a rapid and opportunistic response to the sudden availability of water is clearly an adaptative advantage to access and utilize more favourable, and newly available, ecological substrates and niches.

Water addition led to a general increase in the relative abundance of ribosomal protein transcripts (Rp:T), interpreted as an increase in cellular activity (Bosdriesz et al., 2015; Bremer and Dennis, 1996). Cellular activity levels returned to basal (control) levels within 7 days, in parallel with the desiccation of the soil samples. These data are consistent with the paradigm that desert ecosystems and their indigenous microbiota are both resilient and water-pulse responsive (e.g., Belnap et al., 2005; Noy-Meir, 1973; Armstrong et al., 2016).

Interestingly, various groups of taxa reached maximum Rp:T values at different times, suggesting a controlled pattern of functionality reminiscent of a stepwise model where ecosystem functions gradually evolve as a function of the duration and intensity of the water pulses (Schwinning and Sala, 2004). The most immediate microbial response was characterized by transcription of genes implicated in the motility apparatus (type IV pili in Alpha-proteobacteria and flagella in Actinobacteria). It has been previously noted that one of the main impacts of water inundation is increased soil particle connectivity, providing access to new niches and solubilized nutrients (Schimel, 2018). Actinobacteria and Alpha-proteobacteria have been reported as the dominant active taxa in desiccated soils (León-Sobrino et al., 2019), possibly uniquely positioned to access new and more favourable niches during periods of inter-connection associated with the water-saturated state.

A significant, but delayed, transcriptional activation was observed in the non-fungal microbial eukaryotes (e.g., protists) and Deltaproteobacteria; i.e., 3 to 7 hours after water addition. The former were mostly characterized by structural gene transcripts from the cytoskeleton, a generic indication of overall cellular activity (cell motility and/or cell division). Up-regulated Deltaproteobacterial transcripts were predominantly derived from the Myxococcales, an order of well-known predatory bacteria (Jurkevitch and Davidov, 2007; Shimkets et al., 2006). The dramatic increase in protist and myxobacterial activity is strongly suggestive of predatory behaviour (Thiery and Kaimer, 2020), possibly triggered by increases in prey abundance (i.e., Actinobacteria and Alphaproteobacteria populations) rather than just by soil rehydration.

### Namib Desert fungal and cyanobacterial soil populations are not activated by water

Among the edaphic taxa that were essentially non-responsive to water addition, we particularly identified the Cyanobacteria and most Fungi, with almost none of their genes being differentially up-regulated at a statistically significant level.

We anticipated that water addition would trigger a significant and rapid increase in primary production markers linked to cyanobacterial and photosynthetic activity, as previously observed in biological soil crusts and hypolithic communities (Angel and Conrad, 2013; Pringault and Garcia-Pichel, 2004; Rajeev et al., 2013; Steven et al., 2018; Warren-Rhodes et al., 2006). The almost complete absence of water input-related activation of cyanobacterial functionality suggests that primary productivity in hyperarid soils may not be driven by cyanobacteria and is consistent with previous observations showing that hypolithons (and maybe other cryptic communities) are the foundation of productivity after rain events in the Namib Desert (Ramond et al., 2018).

### Desert edaphic viral communities change over time after a wetting event

The extent to which microbial communities in desert soils are influenced by phage remains largely unexplored (Fancello et al., 2013; Zablocki et al., 2016; 2017). This study provides one of the first temporal assessments of active phage in a desert edaphic environment (Zablocki et al., 2016). In parallel with the rapid changes in bacterial metatranscriptomic patterns, the phage population responded within 10 min after water addition, followed by a sharp decrease and a secondary increase broadly correlating with the ‘late response’ fraction of the soil bacterial community.

We propose two possible mechanisms which could lead to the bimodal response of viral communities in wetted soils: a direct response to increased predation and/or a *kill-the-winner* model as observed in aquatic ecosystems (e.g., Shapiro et al., 2010). Microbial taxa in a ‘biologically hostile’ environment may be forced to allocate resources to energetically expensive defence mechanisms, potentially resulting in lowered defence against phage and elevated susceptibility to infection (Chen and Williams, 2012; Friman and Buckling, 2013; Örmälä-Odegrip et al., 2015). The bacterial host’s capacity to maintain the ‘arms race’ dynamic (i.e., co-escalation of host resistance and parasite infectivity) may also be degraded in the presence of predators (Friman and Buckling, 2013).

We note that most of the viral sequences identified in this study were unclassified suggesting, as previously observed (Scola et al., 2018), that hot desert soils may harbour a substantial and potentially novel phage genomic and taxonomic diversity.

### Water pulses shift microbial C, N and P nutrient utilization patterns

It was anticipated that water addition would trigger a significant increase in primary productivity, since photosynthetic processes are highly sensitive to water activity (Brock, 1975; Steven et al., 2018; Warren-Rhodes et al., 2006). However, markers for photosynthetic and chemoautotrophic carbon fixation (the latter being active in desiccated periods (León-Sobrino et al., 2019; Sghaier et al., 2016)), were significantly suppressed. This suggests that open soil communities may be dependent on carbon input from alternative sources, such as sporadic vegetation growth and/or productive cryptic niches such as hypolithons and biological soil crusts (Armstrong et al., 2016; Ramond et al., 2018). Nitrogen cycling genes, particularly those involved in inorganic nitrogen acquisition (i.e., nitrate, via nitrate reductases), were also downregulated after water addition. This was also surprising as active N mineralization and N loss processes are often increased in arid soils in response to rainfall (Austin et al., 2004), and desert soils are particularly rich in nitrate (Graham et al., 2008; Walwoord et al., 2003). However, a metabolic switch to nitrogen acquisition from organic substrates was strongly suggested by the upregulation of peptide transporter genes, mirroring the situation observed for carbon acquisition. The transient reduction in autotrophic C and N fixation after watering may be explained in terms of energy efficiency, where the sudden availability of ‘energy-rich’ substrates provides a favoured heterotrophic resource over energetically expensive autotrophic processes (Fuchs, 2011).

The addition of water triggered an upregulation of genes involved in inorganic phosphate transport and a downregulation of those implicated in organic phosphonate acquisition. We speculate that the solubilization of inorganic phosphate from soil particles displaces phosphonates as the preferred P source (Schowanek and Verstraete, 1990). Noticeable exceptions were the Alpha-proteobacterial taxa, which apparently favour organic G3P as a preferred P source, both in desiccated soils and after wetting (León-Sobrino et al., 2019).

### A conceptual response model of desert soil edaphic microbial communities to water

From a composite analysis of our metatranscriptomic data, we propose a rainfall response model for desert soil microbiomes (Figure 8). Immediately after soil wetting (≤ 10 min), some bacterial taxa (particularly Actinobacteria and Alpha-proteobacteria, that show significant chemoautotrophic capacity in dry soils (León-Sobrino et al., 2019)) reduce autotrophic carbon fixation activities, activate cellular uptake mechanisms and engage in dispersal using both natatory (flagella) and gliding (type IV pili) mechanisms, presumably in order to colonize new niches and access new substrate resource pools. There is a concomitant increase in active phage particles.

**Figure 8.**
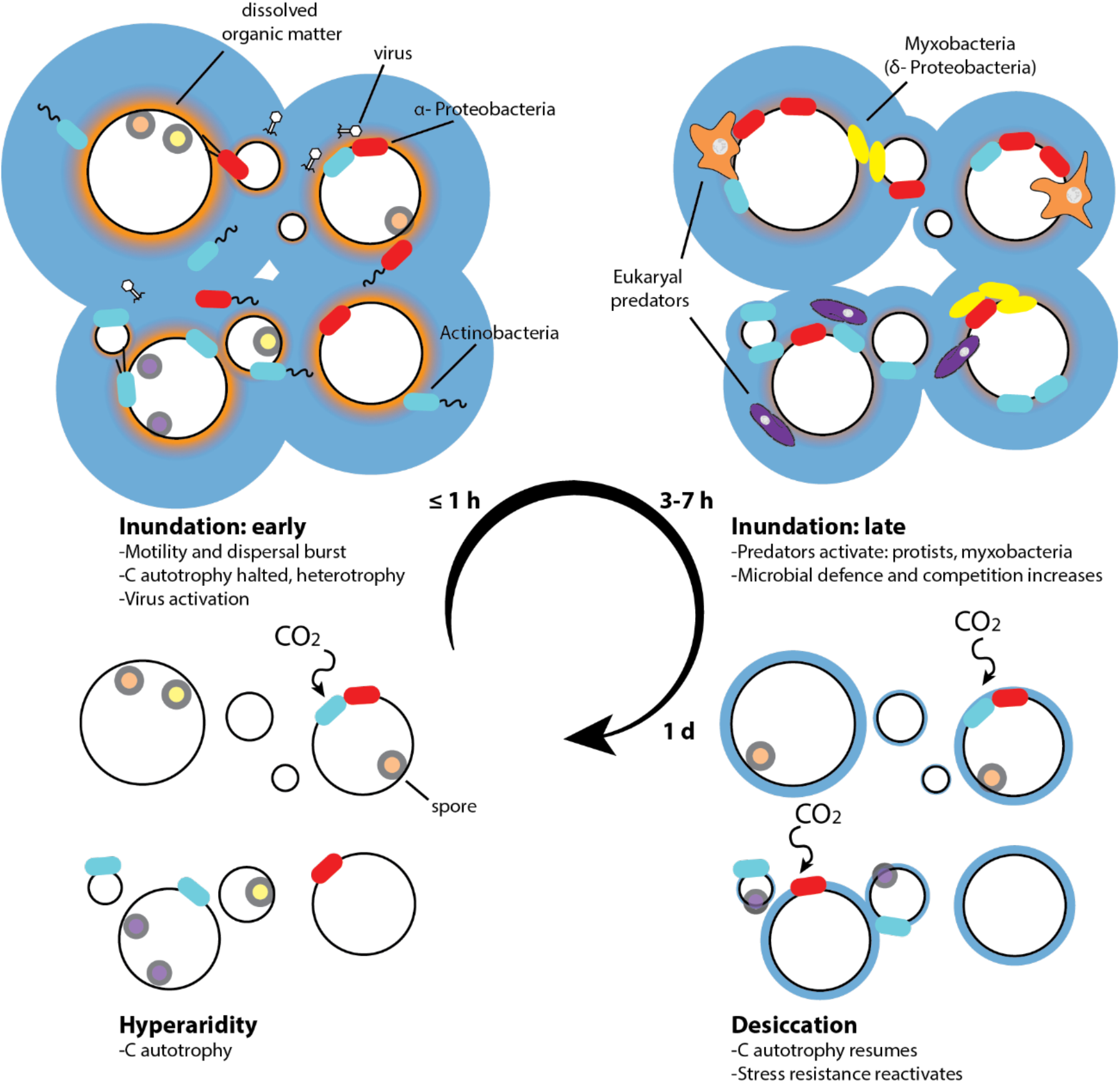
Response model of microbial communities to water events in hyperarid desert soils.

Following this rapid dispersal burst, approximately 3 to 7 hours after water addition predatory and saprophytic microbial taxa are activates. These predators include eukaryotes, especially ciliates (Oligohymenophorea class), Dictyostellid amoebae, Delta-proteobacteria (myxobacteria) and Bacteroidetes (Cytophagia class). Simultaneously, and presumably in response to the activation of predators, several bacterial groups upregulate the transcription of defensive systems, most notably T6SS. A second wave of viral infections appears to follow the emergence of eukaryotic predators.

In the final stage of the wetting-drying cycle (approx. 7 days after water addition), when soils are effectively dehydrated to pre-watering levels, autotrophic carbon fixation processes and nitrogen cycling are reactivated in the bacterial community, along with certain desiccation stress resistance mechanisms, most especially HSP20.

## Conclusion

In this 7 day *in situ* metatranscriptomics study, we have observed the dominant microbial processes induced by ‘rainfall’ on hyperarid soils and successfully followed a complete cycle of microbial activation and subsequent inactivation as soils and their microbial communities returned to their basal desiccated state and transcriptional status. We show strong evidence of short-term temporal succession and, by implication, tightly regulated processes in desert edaphic communities after water addition, which favour heterotrophy over autotrophy. Furthermore, we suggest that dispersal-predation cycles, in which both phage and predators play active roles, are important driving forces in shaping community composition within a wetting-drying regime in hyper-arid desert soils.

## Declarations

## Acknowledgements

The authors wish to thank the National Research Foundation of South Africa and the University of Pretoria for financial support, and the staff and students of the Gobabeb-Namib Research Institute for field support.

## Data availability

The dataset supporting the conclusions of this article is available in the ArrayExpress repository, https://www.ebi.ac.uk/arrayexpress/ (accession no. E-MTAB-9439). Reference metatranscriptome assembly and annotations can be accessed at the IMG/M repository, https://img.jgi.doe.gov/ (GOLD Analysis Project Id Ga0326365)

## Author contributions

C. L.-S., J.-B. R. and D. A. C. conceived the experiment. C. L.-S., J.-B. R. and R. K. performed the sample collection. C. L.-S. performed all experimental work. C. L-S. and C. C. performed the microbial and viral computational data analysis, respectively. C. L.-S., C. C., J.-B. R. and D. A. C. participated in the interpretation of results and writing of the manuscript. G. M.-K. provided logistical support and field advice in the Namib Desert.

