## Supplementary materials for "Temporal dynamics of microbial transcription in wetted hyperarid desert soils"

### *SUPPLEMENTARY TABLES*

| Sector | pH | Na mg/l | K mg/l | Ca mg/l | Mg mg/l | $NO_3^-$ mg/l | $NH_4^+$ mg/l | Cl mg/l | EG mS/m | P mg/l | C % | Sand % | Clay % | Silt % | >1000 µm % | >500 µm % | >250 µm % | >100 µm % | >53 µm % | <53 µm % |
|---|---|---|---|---|---|---|---|---|---|---|---|---|---|---|---|---|---|---|---|---|
| 10 min C2 | 7.5 | 5.26 | 2 | 2.6 | 1.03 | 0.3 | 0.1 | 9.7 | 8 | 0.19 | 0.03 | 95 | 6 | - | 4.80 | 5.15 | 5.90 | 45.47 | 33.19 | 2.63 |
| 10 min C3 | 7.5 | 5.67 | 2.26 | 3.86 | 1.13 | 0.9 | 1.4 | 11.9 | 9 | 0.2 | 0.06 | 91 | 6 | 3 | 2.48 | 3.64 | 4.43 | 40.21 | 40.47 | 5.28 |
| 10 min W1 | 7.4 | 5.44 | 2.84 | 5.95 | 1.44 | 0.1 | 0.1 | 11.5 | 11 | 0.25 | 0.09 | 91 | 8 | 1 | 4.34 | 5.00 | 4.62 | 41.10 | 35.72 | 2.72 |
| 10 min W2 | 7.5 | 5.15 | 3.04 | 4.17 | 1.37 | 0.7 | 0.7 | 10.8 | 9 | 0.29 | 0.03 | 90 | 6 | 4 | 3.80 | 4.59 | 5.13 | 43.87 | 33.07 | 4.84 |
| 1 h C1 | 7.4 | 5.04 | 2.48 | 2.88 | 1.15 | 0.9 | 2.3 | 12.6 | 8 | 0.2 | 0.03 | 93 | 6 | 1 | 3.44 | 4.05 | 4.71 | 41.70 | 39.51 | 3.23 |
| 1 h C2 | 7.3 | 6.06 | 2.46 | 5.2 | 1.31 | 0.2 | 0.1 | 12.2 | 10 | 0.18 | 0.05 | 93 | 6 | 1 | 3.07 | 3.76 | 5.58 | 47.52 | 33.20 | 4.58 |
| 1 h W2 | 7.1 | 5.89 | 3.23 | 6.73 | 1.45 | 0 | 1.6 | 11.5 | 12 | 0.26 | 0.05 | 91 | 6 | 3 | 3.42 | 3.99 | 4.44 | 42.05 | 37.43 | 5.43 |
| 1 h W3 | 7.2 | 5.08 | 2.97 | 7.44 | 1.51 | 2.6 | 0.6 | 11.6 | 12 | 0.23 | 0.06 | 92 | 6 | 2 | 3.49 | 4.10 | 4.87 | 43.30 | 36.72 | 5.01 |
| 3 h C1 | 7.5 | 5.86 | 2.34 | 3.88 | 1.33 | 0.6 | 1.1 | 11.5 | 9 | 0.22 | 0.08 | 93 | 5 | 2 | 3.49 | 3.95 | 4.67 | 43.15 | 37.59 | 3.97 |
| 3 h C2 | 7.5 | 5.53 | 2.32 | 4.48 | 1.27 | 0.6 | 0.7 | 14.4 | 8 | 0.21 | 0.07 | 92 | 6 | 2 | 3.20 | 3.95 | 4.98 | 45.35 | 34.75 | 4.46 |
| 3 h W1 | 7.3 | 6.12 | 3.12 | 5.39 | 1.48 | 0.8 | 0.1 | 14.4 | 11 | 0.33 | 0.02 | 90 | 6 | 4 | 3.52 | 4.72 | 4.70 | 36.69 | 40.67 | 5.18 |
| 3 h W2 | 7.3 | 5.29 | 2.69 | 4.77 | 1.19 | 0.1 | 0.2 | 10.8 | 10 | 0.29 | 0.03 | 93 | 6 | 1 | 3.44 | 4.27 | 5.13 | 45.48 | 34.38 | 4.31 |
| 7 h C2 | 7.5 | 5.55 | 2.81 | 3.85 | 1.2 | 0.8 | 0.8 | 13 | 10 | 0.19 | 0.05 | 92 | 6 | 2 | 2.55 | 3.27 | 4.23 | 43.11 | 39.05 | 4.73 |
| 7 h C3 | 7.6 | 5.4 | 2.19 | 4.43 | 1.19 | 0.8 | 0.8 | 11.5 | 9 | 0.18 | 0.02 | 94 | 7 | - | 3.99 | 4.01 | 4.74 | 47.69 | 33.81 | 3.56 |
| 7 h W2 | 7.7 | 5.67 | 2.94 | 5.01 | 1.24 | 1.7 | 0.6 | 10.4 | 11 | 0.27 | 0.05 | 91 | 6 | 3 | 3.44 | 4.57 | 4.94 | 45.09 | 33.37 | 4.88 |
| 7 h W3 | 7.5 | 4.87 | 2.7 | 5.41 | 1.31 | 1.1 | 0.6 | 11.2 | 10 | 0.3 | 0.06 | 93 | 6 | 1 | 6.36 | 7.17 | 6.33 | 42.48 | 30.55 | 3.42 |
| 1 d C2 | 7.1 | 5.55 | 2.56 | 3.39 | 1.34 | 1.2 | 0.7 | 9.7 | 8 | 0.25 | 0.04 | 92 | 6 | 2 | 2.64 | 3.36 | 4.44 | 44.30 | 37.51 | 5.13 |
| 1 d C3 | 7.1 | 5.41 | 2.45 | 4.37 | 1.36 | 1 | 1.7 | 12.2 | 8 | 0.22 | 0.02 | 93 | 4 | 3 | 2.82 | 4.69 | 5.82 | 46.84 | 32.72 | 4.43 |
| 1 d W2 | 7.4 | 4.03 | 2.54 | 2.82 | 1.68 | 1 | 1 | 7.2 | 6 | 0.34 | 0.03 | 92 | 6 | 2 | 2.94 | 3.42 | 4.45 | 50.55 | 30.72 | 4.56 |
| 1 d W3 | 7.2 | 4.19 | 2.91 | 2.96 | 1.59 | 0.3 | 1.2 | 8.6 | 7 | 0.42 | 0.04 | 90 | 7 | 3 | 5.04 | 5.68 | 5.64 | 37.62 | 36.17 | 4.63 |
| 7 d C2 | 7.3 | 4.6 | 2.36 | 2.85 | 1.27 | 0.9 | 1.7 | 6.8 | 7 | 0.23 | 0.03 | 93 | 7 | 0 | 2.37 | 3.91 | 5.56 | 46.96 | 34.08 | 3.78 |
| 7 d C3 | 7.5 | 4.98 | 2.49 | 3.61 | 1.27 | 1 | 1.6 | 11.2 | 8 | 0.17 | 0.04 | 92 | 5 | 3 | 2.49 | 3.53 | 4.92 | 44.58 | 36.66 | 4.24 |
| 7 d W2 | 7.5 | 4.23 | 2.04 | 2.66 | 1.19 | 0.6 | 0.9 | 6.8 | 6 | 0.2 | 0.05 | 92 | 6 | 2 | 3.33 | 3.97 | 4.87 | 43.99 | 35.84 | 4.11 |
| 7 d W3 | 8 | 4.3 | 2.56 | 3.24 | 1.46 | 0.7 | 1.3 | 8.6 | 7 | 0.21 | 0.02 | 87 | 8 | 5 | 2.47 | 3.43 | 4.08 | 40.09 | 36.88 | 6.07 |

**Table S1**: Soil composition of samples selected for transcriptome sequencing. C and W in sample identifiers refer to Control (dry) and Watered plots, respectively. All soil properties were determined from particles >2 mm.

| | | **Dry** | | | | | | **Watered** | | | | | | **Dry 7-hour ave.** | **Watered 7-hour ave.** |
|---|---|---|---|---|---|---|---|---|---|---|---|---|---|---|---|
| **Class** | **Phylum** | **10 min** | **1 h** | **3 h** | **7 h** | **1 d** | **7 d** | **10 min** | **1 h** | **3 h** | **7 h** | **1 d** | **7 d** | | |
| Actinobacteria | | 29.66 | 21.82 | 24.11 | 19.60 | 21.99 | 21.24 | 29.51 | 19.53 | 19.52 | 22.18 | 18.83 | 20.28 | 23.80 | 22.68 |
| Rubrobacteria | | 16.90 | 17.73 | 15.71 | 15.81 | 14.94 | 11.68 | 10.97 | 10.57 | 10.76 | 9.04 | 9.30 | 12.36 | 16.54 | 10.34 |
| Thermoleophilia | Actinobacteria | 4.31 | 3.21 | 3.57 | 3.03 | 2.80 | 3.02 | 5.54 | 4.87 | 4.05 | 2.90 | 3.55 | 4.44 | 3.53 | 4.34 |
| Other | | 0.66 | 0.72 | 0.61 | 0.81 | 0.86 | 1.86 | 0.94 | 0.91 | 1.03 | 0.66 | 0.66 | 0.67 | 0.70 | 0.88 |
| Total | | 51.53 | 43.47 | 44.01 | 39.25 | 40.59 | 37.79 | 46.96 | 35.87 | 35.35 | 34.78 | 32.34 | 37.76 | 44.56 | 38.24 |
| Alphaproteobacteria | | 9.38 | 10.00 | 12.07 | 9.87 | 9.91 | 9.84 | 12.34 | 9.90 | 13.23 | 12.52 | 8.02 | 5.66 | 10.33 | 12.00 |
| Deltaproteobacteria | | 1.43 | 1.46 | 1.81 | 1.43 | 2.07 | 3.75 | 3.02 | 3.48 | 5.92 | 6.16 | 3.73 | 1.63 | 1.54 | 4.64 |
| Gammaproteobacteria | Proteobacteria | 1.24 | 1.31 | 1.20 | 1.23 | 1.18 | 1.52 | 1.13 | 1.45 | 1.49 | 1.44 | 1.54 | 1.87 | 1.24 | 1.38 |
| Betaproteobacteria | | 1.61 | 1.74 | 1.72 | 1.61 | 1.55 | 2.29 | 1.18 | 1.14 | 1.18 | 1.42 | 1.64 | 2.46 | 1.67 | 1.23 |
| Other | | 0.26 | 0.30 | 0.36 | 0.34 | 0.27 | 0.29 | 0.18 | 0.20 | 0.16 | 0.22 | 0.19 | 0.42 | 0.32 | 0.19 |
| Total | | 13.92 | 14.81 | 17.15 | 14.49 | 14.97 | 17.69 | 17.84 | 16.16 | 21.97 | 21.77 | 15.12 | 12.03 | 15.09 | 19.44 |
| Nitrososphaeria | | 8.62 | 13.72 | 9.23 | 16.33 | 14.73 | 13.63 | 4.34 | 15.40 | 10.36 | 11.04 | 23.90 | 22.27 | 11.98 | 10.28 |
| Other | Thaumarchaeota | 0.15 | 0.24 | 0.15 | 0.38 | 0.21 | 0.26 | 0.08 | 0.25 | 0.14 | 0.16 | 0.45 | 0.40 | 0.23 | 0.16 |
| Total | | 8.76 | 13.96 | 9.38 | 16.71 | 14.94 | 13.89 | 4.42 | 15.65 | 10.50 | 11.20 | 24.36 | 22.67 | 12.20 | 10.44 |
| Chloroflexia | | 4.30 | 4.32 | 6.24 | 3.95 | 4.58 | 2.87 | 4.37 | 4.06 | 3.87 | 2.42 | 3.12 | 2.39 | 4.70 | 3.68 |
| unclass. Chloroflexi | Chloroflexi | 1.20 | 1.15 | 1.61 | 0.95 | 1.08 | 1.94 | 0.46 | 0.65 | 0.47 | 0.39 | 0.93 | 1.66 | 1.23 | 0.49 |
| Other | | 4.77 | 4.57 | 5.81 | 4.04 | 3.60 | 3.65 | 3.27 | 3.01 | 2.91 | 1.91 | 2.99 | 4.19 | 4.80 | 2.78 |
| Total | | 10.27 | 10.05 | 13.66 | 8.94 | 9.26 | 8.46 | 8.09 | 7.72 | 7.26 | 4.72 | 7.04 | 8.24 | 10.73 | 6.95 |
| Bacilli | | 1.89 | 2.45 | 2.06 | 3.05 | 3.09 | 5.10 | 1.69 | 2.53 | 2.85 | 2.27 | 2.27 | 2.37 | 2.37 | 2.34 |
| Clostridia | Firmicutes | 1.01 | 1.27 | 0.97 | 1.01 | 1.34 | 3.59 | 1.46 | 1.62 | 1.53 | 1.11 | 1.22 | 1.36 | 1.06 | 1.43 |
| Other | | 0.17 | 0.13 | 0.13 | 0.12 | 0.09 | 0.13 | 0.20 | 0.20 | 0.19 | 0.13 | 0.17 | 0.16 | 0.14 | 0.18 |
| Total | | 3.07 | 3.85 | 3.16 | 4.19 | 4.53 | 8.81 | 3.35 | 4.35 | 4.57 | 3.51 | 3.65 | 3.89 | 3.57 | 3.94 |

| | | | | | | | | | | | | | | | |
|---|---|---|---|---|---|---|---|---|---|---|---|---|---|---|---|
| Cytophagia | | 0.75 | 0.76 | 0.56 | 1.26 | 1.15 | 0.62 | 1.70 | 1.78 | 1.41 | 3.22 | 1.11 | 0.77 | 0.83 | 2.03 |
| Sphingobacteriia | | 0.83 | 0.81 | 0.64 | 1.97 | 1.70 | 0.73 | 0.95 | 0.72 | 0.64 | 1.28 | 0.48 | 0.42 | 1.06 | 0.90 |
| Chitinophagia | Bacteroidetes | 0.22 | 0.26 | 0.22 | 0.63 | 0.39 | 0.26 | 0.44 | 0.61 | 0.61 | 1.05 | 0.41 | 0.21 | 0.34 | 0.68 |
| Other | | 0.22 | 0.28 | 0.26 | 0.30 | 0.26 | 0.22 | 0.47 | 0.59 | 0.51 | 0.58 | 0.53 | 0.27 | 0.26 | 0.54 |
| Total | | 2.02 | 2.11 | 1.69 | 4.16 | 3.49 | 1.83 | 3.57 | 3.70 | 3.17 | 6.14 | 2.52 | 1.67 | 2.50 | 4.15 |
| Planctomycetia | | 1.66 | 1.76 | 2.20 | 1.59 | 1.34 | 1.41 | 2.53 | 1.85 | 1.65 | 1.76 | 1.56 | 1.97 | 1.81 | 1.95 |
| Other | Planctomycetes | 0.10 | 0.08 | 0.08 | 0.07 | 0.09 | 0.06 | 0.43 | 0.47 | 0.46 | 0.43 | 0.29 | 0.07 | 0.08 | 0.45 |
| Total | | 1.76 | 1.85 | 2.29 | 1.67 | 1.43 | 1.47 | 2.96 | 2.32 | 2.11 | 2.19 | 1.85 | 2.05 | 1.89 | 2.39 |
| unclass. Cyanobacteria | | 1.54 | 1.59 | 1.57 | 1.63 | 1.47 | 1.55 | 0.99 | 0.99 | 1.07 | 0.94 | 1.41 | 2.17 | 1.58 | 1.00 |
| Other | Cyanobacteria | 0.06 | 0.06 | 0.07 | 0.07 | 0.06 | 0.10 | 0.05 | 0.03 | 0.04 | 0.02 | 0.05 | 0.08 | 0.06 | 0.03 |
| Total | | 1.60 | 1.65 | 1.65 | 1.70 | 1.53 | 1.64 | 1.03 | 1.02 | 1.11 | 0.96 | 1.45 | 2.25 | 1.65 | 1.03 |
| Blastocatellia | | 0.42 | 0.30 | 0.44 | 0.60 | 0.30 | 0.36 | 0.63 | 1.42 | 1.12 | 0.41 | 0.70 | 0.37 | 0.44 | 0.89 |
| Other | Acidobacteria | 0.64 | 0.62 | 0.65 | 0.69 | 0.57 | 0.93 | 0.71 | 0.92 | 0.91 | 0.74 | 0.93 | 0.79 | 0.65 | 0.82 |
| Total | | 1.06 | 0.92 | 1.09 | 1.29 | 0.87 | 1.29 | 1.34 | 2.33 | 2.03 | 1.14 | 1.64 | 1.16 | 1.09 | 1.71 |
| Pezizomycetes | | 0.20 | 0.29 | 0.05 | 0.43 | 0.48 | 0.14 | 1.70 | 0.74 | 0.23 | 0.76 | 0.24 | 0.24 | 0.24 | 0.86 |
| Other | Ascomycota | 0.40 | 0.48 | 0.39 | 0.52 | 1.71 | 0.36 | 0.87 | 0.40 | 0.35 | 0.66 | 0.52 | 0.65 | 0.44 | 0.57 |
| Total | | 0.60 | 0.77 | 0.44 | 0.95 | 2.19 | 0.50 | 2.57 | 1.13 | 0.58 | 1.42 | 0.76 | 0.89 | 0.69 | 1.43 |
| Oligohymenophorea | | 0.10 | 0.18 | 0.11 | 0.50 | 0.34 | 0.27 | 0.42 | 1.81 | 2.62 | 2.43 | 0.49 | 0.13 | 0.22 | 1.82 |
| unclass. Eukaryota | unclass. Eukaryota | 0.16 | 0.26 | 0.13 | 0.23 | 0.25 | 0.12 | 0.21 | 0.43 | 0.68 | 1.66 | 0.87 | 0.14 | 0.19 | 0.74 |
| Other | | 0.03 | 0.08 | 0.10 | 0.18 | 0.07 | 0.08 | 0.04 | 0.04 | 0.03 | 0.34 | 0.04 | 0.10 | 0.10 | 0.11 |
| Total | | 0.29 | 0.53 | 0.34 | 0.90 | 0.66 | 0.46 | 0.67 | 2.28 | 3.32 | 4.43 | 1.40 | 0.37 | 0.51 | 2.67 |
| Gemmatimonadetes | Gemmatimonadetes | 0.30 | 0.23 | 0.20 | 0.24 | 0.22 | 0.21 | 1.65 | 1.33 | 1.18 | 1.29 | 1.05 | 0.23 | 0.24 | 1.36 |
| Total | | 0.30 | 0.23 | 0.20 | 0.24 | 0.22 | 0.21 | 1.65 | 1.33 | 1.18 | 1.29 | 1.05 | 0.23 | 0.24 | 1.36 |
| **REST** | | 4.82 | 5.79 | 4.96 | 5.53 | 5.34 | 5.95 | 5.55 | 6.12 | 6.86 | 6.44 | 6.83 | 6.80 | 5.27 | 6.24 |
| **Classified TPM** | | 41.36 | 37.52 | 39.64 | 36.72 | 37.16 | 33.50 | 41.02 | 39.00 | 38.29 | 36.61 | 36.84 | 30.52 | | |

**Table S2**: Taxonomic classification of gene transcripts as a percentage of all classified transcripts (TPM). Only phyla with transcript abundances >1% on the 7-hour averages (dry or watered) are shown. “Classified TPM” percentages indicated in the bottom row are calculated from the total.

### *SUPPLEMENTARY FIGURES*

| | | | | | | |
|---|---|---|---|---|---|---|
| 7 h 1 | 2 h 3 | 3 d 2 | 10 min 3 | 1 d 2 | - | 4 d 3 |
| 3 d 3 | 7 d 1 | - | 1 d 1 | 1 h 1 | - | - |
| 1 mo 1 | 7 d 3 | - | - | 2 h 2 | 10 min 1 | 2 d 1 |
| 2 d 2 | - | - | 2 d 3 | 2 mo 2 | 7 h 2 | - |
| 1 d 3 | - | 1 mo 3 | - | 3 h 3 | 2 mo 3 | 1 h 2 |
| 2 h 1 | 3 d 1 | 7 h 3 | 4 d 1 | 4 d 2 | 10 min 2 | 2 mo 1 |
| 1 h 3 | - | - | 3 h 2 | 7 d 2 | 1 mo 2 | 3 h 1 |

**Figure S1**: Spatial distribution of the sampled soils within the sampling plots. The same positions were used for the dry and watered plots. The times for which samples were sequenced are coloured. Refer to Table S1 for more details on the sequenced sectors.

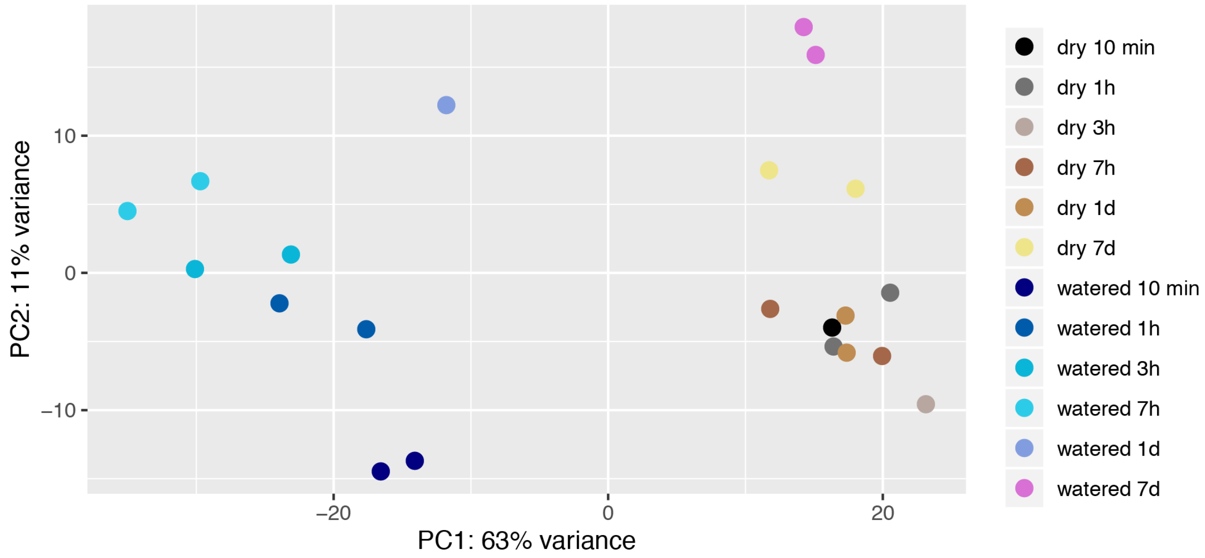


**Figure S2**: principal components analysis of transcriptome reads classified along combined taxonomic (class) and functional (KO) categories. Computed by *DESeq2*.

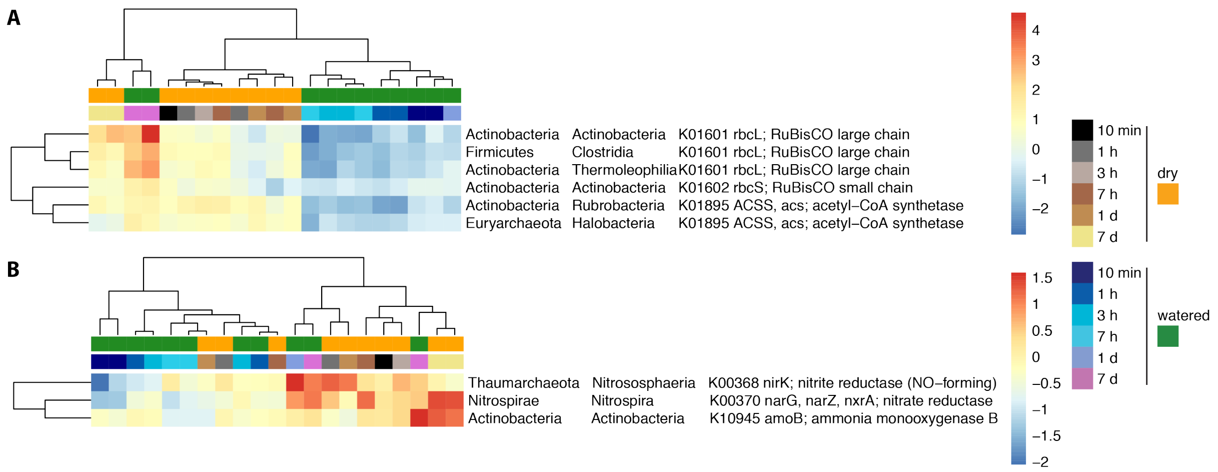


**Figure S3**: A) Carboxylase genes from carbon fixation pathways and B) nitrogen cycling enzyme genes significantly changing after watering. Transcript data was aggregated along combined taxonomic (class) and functional (KEGG Orthologs) groups for the analysis. Values were normalized using the Variance Stabilizing Transformation (*DESeq* R package). Rows and columns were clustered using *hclust*.
